# Delayed antigen-specific CD4^+^ T-cell induction correlates with impaired immune responses to SARS-COV-2 mRNA vaccination in the elderly

**DOI:** 10.1101/2022.05.10.490700

**Authors:** Norihide Jo, Yu Hidaka, Osamu Kikuchi, Masaru Fukahori, Takeshi Sawada, Masahiko Aoki, Masaki Yamamoto, Miki Nagao, Satoshi Morita, Takako E Nakajima, Manabu Muto, Yoko Hamazaki

**Affiliations:** Department of Life Science Frontiers, Center for iPS Cell Research and Application (CiRA), Kyoto University; Sakyo-Ku, Kyoto 606-8507, Japan; Alliance Laboratory for Advanced Medical Research, Graduate school of Medicine, Kyoto University; Sakyo-Ku, Kyoto 606-8507, Japan; Department of Biomedical Statistics and Bioinformatics, Graduate School of Medicine, Kyoto University; Sakyo-Ku, Kyoto 606-8507, Japan; Department of Therapeutic Oncology, Graduate School of Medicine, Kyoto University; Sakyo-Ku, Kyoto 606-8507, Japan; Clinical Bio-Resource Center, Kyoto University Hospital; Sakyo-Ku, Kyoto 606-8507, Japan; Department of Early Clinical Development, Graduate school of Medicine, Kyoto University; Sakyo-Ku, Kyoto 606-8507, Japan; Kyoto Innovation Center for Next Generation Clinical Trials and iPS Cell Therapy (Ki-CONNECT), Kyoto University Hospital; Sakyo-Ku, Kyoto 606-8507, Japan; Department of Clinical Laboratory Medicine, Graduate School of Medicine, Kyoto University; Sakyo-Ku, Kyoto 606-8507, Japan; Laboratory of Immunobiology, Graduate school of Medicine, Kyoto University; Sakyo-Ku, Kyoto 606-8507, Japan

## Abstract

Despite the clinical efficacy of coronavirus disease 2019 mRNA vaccines, the elderly demonstrate lower IgG levels and neutralizing titers and a higher risk of severe diseases. CD4^+^ T cells play a central role in regulating antigen-specific antibody and CD8^+^ T-cell responses; however, because their composition and functionality change significantly with age, relationships between age-associated defects in T cells and the immunogenicity of or reactogenicity to mRNA vaccines are unclear. Using a vaccine cohort (*n* = 216), we found that the elderly (aged ≥65 years) showed delayed induction and early contraction of vaccine-specific CD4^+^ T cells, and that the compromised C–X–C motif chemokine receptor 3^+^ circulating T follicular helper cell response after the first dose was associated with the lower IgG levels. Additionally, the elderly experienced significantly fewer systemic adverse effects (AEs) after the second dose, with those exhibiting few AEs showing lower cytokine^+^ CD4^+^ T cells after the first dose and lower antibody levels after the second dose. Furthermore, T helper 1 cells in the elderly expressed higher levels of programmed cell death protein-1, a negative regulator of the T-cell response, which was associated with less production of vaccine-specific CD4^+^ T cells and impaired CD8^+^ T-cell expansion. Thus, efficient induction of vaccine-specific effector/memory CD4^+^ T cells after the first dose may trigger robust cytokine production after the second dose, leading to effective vaccine responses and higher systemic reactogenicity. These results suggested that an enhanced CD4^+^ T-cell response after the first dose is key to improved vaccination efficacy in the elderly.

**One Sentence Summary:** We compared immunogenicity and reactogenicity to COVID-19 mRNA vaccine in 107 adults (aged <65 years) and 109 elderly (aged ≥65) individuals.

## INTRODUCTION

Advanced age is the strongest risk factor for severe symptoms and death with coronavirus disease 2019 (COVID-19) (*1-3*), and this may be largely due to the age-associated decline in immune competence. Aging leads to various changes that affect nearly every component of the immune system. T cells are immune cells that belong to the adaptive immune system and play a central role in antigen-specific antiviral immune responses, including antibody response and cytotoxicity against virus-infected cells (*4*). Despite their critical roles, the production of new T cells starts to decrease in early life stages due to thymic involution and undergoes various qualitative and compositional changes and functional dysregulations with age (*5-10*). Thus, the elderly are strongly recommended to receive vaccines; however, the benefits and efficacy of vaccination are limited, mainly because of the decreased effectiveness of acquired immunity (*11*). Indeed, vaccines against influenza or pneumococcal diseases that comprise inactivated virus or subunit proteins can mitigate the symptoms but do not induce protective immunity in the elderly (*12, 13*).

The newly developed severe acute respiratory syndrome coronavirus 2 (SARS-CoV-2) mRNA vaccines are highly effective at preventing severe illness, as well as infection, at ∼95% efficacy 7 days after the second dose, even in subjects aged >75 years (*14*). However, vaccine-specific IgG levels and neutralizing antibody titers are significantly lower in the elderly for up to 6 months after the second dose (*15-17*). Thus, the elderly are still at increased risk of reduced vaccine effectiveness several months after vaccination and vulnerable to severe disease, hospitalization, and death (*16*). A detailed immunological study revealed that the mRNA vaccines elicit strong T helper (Th)1- and T follicular helper (Tfh)-cell responses (*18, 19*). Importantly, individuals >80 years of age showed lower IgG titer and serum neutralization activity as well as fewer cytokine^+^ CD4^+^ T cells after vaccination (*15*). However, the detailed trajectory of T-cell responses and how Th1 and Tfh cell responses are affected in the elderly remains to be investigated. Considering the importance of T cells in vaccine responses, elucidating age-associated differences in T-cell responses to mRNA vaccines is fundamental.

Noticeable and severe adverse effects (AEs) are another characteristic of mRNA vaccines (*20*). Notably, AEs have been more frequently observed after the second dose and with higher severity than after the first dose (*14*), which strongly suggested that AEs are a consequence of immunological memory induced by the adaptive immune system. However, previous reports did not reach a consensus concerning the association between AEs and vaccine-induced immune reactions, such as antibody production, likely due to the small cohorts used in the studies and variations in how AEs were defined (*21-27*). Moreover, most studies examined the associations of AEs with antibody levels and/or neutralizing activity but not with T-cell responses, which can result in the production of cytokines and thus cause systemic effects.

In this study, we investigated these key questions by comparing the vaccine-specific Th1- and Tfh-cell responses to two doses of mRNA vaccine between healthy adults and elderly groups in a reasonably large Japanese cohort (*n* = 216) over a period of 3 months post vaccination, including the priming and contraction phases. Furthermore, we explored the associations of vaccine-specific T-cell responses with AEs, as well as the possible mechanism generating the heterogeneity of the immune responses. The results provide insights into understanding the age-related and individual differences in the effectiveness of mRNA vaccines and may be relevant for future vaccine strategies, especially for the highly vulnerable elderly population.

## RESULTS

### Delayed induction and early contraction of vaccine-specific CD4^+^ T cells in the elderly

We studied 216 SARS-CoV-2-naïve Japanese donors comprising adults (aged <65 years, median age: 43 years; *n* = 107) and the elderly (aged ≥65 years, median age: 71 years; *n* = 109) who met the eligibility criteria (see Study design in the Materials and Methods section), having received two doses of BNT162b2 vaccine within around 3-week intervals [Median: 21.0 days (range: 19.0 to 30.0 days)], and were successfully followed up until 3 months after the first dose (Fig. S1A, B and Table 1). Blood samples were obtained before the vaccination [Pre; median: −14 days (range:−29 to 0 days)], 2 weeks after the first dose [Post1; median: 11 days (range: 6−21 days)], 2 weeks after the second dose [Post2; median: 34 days (range: 30−39 days)], and 3 months after the first dose [3 mo; median: 93days (range: 77−104 days)] (Fig. S1A). Donors were also followed up for medical conditions at each study visit. None of the donors tested positive for anti-SARS-CoV-2 nucleocapsid (N) protein IgM/IgG, which reflects the history of COVID-19 during the follow-up period (Table 1).

**Table 1.** Participant characteristics. N: number of subjects in the specified group. This value is the denominator of the percentage calculations. n: Number of subjects with specified characteristics.

|  |  |  | < 65 years<br>N = 107 | ≥ 65 years<br>N = 109 |
| --- | --- | --- | --- | --- |
| Age (year) | - Median (Min, Max) |  | 43 ( 23, 63 ) | 71 ( 65, 81 ) |
| Sex | - n (%) | Male | 43 ( 40.2 % ) | 56 ( 51.4 % ) |
|  |  | Female | 64 ( 59.8 % ) | 53 ( 48.6 % ) |
| Cytomegalovirus IgG | - n (%) | Negative | 34 ( 31.8 % ) | 9 ( 8.3 % ) |
|  |  | Positive | 73 ( 68.2 % ) | 100 ( 91.7 % ) |
| SARS-CoV-2 N IgM/IgG | - n (%) | Negative | 107 ( 100 % ) | 109 ( 100 % ) |
|  |  | Positive | 0 ( 0 % ) | 0 ( 0 % ) |

To quantify and characterize vaccine-specific T-cell response, we utilized activation-induced marker (AIM) and intracellular cytokine staining (ICS) assays. Peripheral blood mononuclear cells (PBMCs) were stimulated with overlapping peptide pools covering the complete sequence of the spike protein of SARS-CoV-2. Markers used for flow cytometric analysis and the gating strategies are shown in Table S1 and Fig. S2 and S3. The total number of CD4^+^ T cells in peripheral blood did not differ between adults and the elderly and remained stable during the study duration in both groups (Fig. 1A). The frequencies and numbers of vaccine-specific AIM^+^ (CD154^+^CD137^+^) CD4^+^ T cells in most donors exhibited a significant increase (median: >10-fold) as compared with the baseline after the first dose, were largely maintained after the second dose, and declined at 3 months (Fig. 1A and B), as reported previously (*28-30*). Vaccine-induced T cells mostly present CCR7^+^CD45RA^−^ central memory (CM) (*30*) and non-senescent (CD28^+^ or CD57^−^) characteristics after the first dose, and this phenotype pattern was maintained until 3 months in both groups (Fig. S4A and B). Optimized t-Distributed Stochastic Neighbor Embedding (opt-SNE), multi-dimension reduction strategy of multicolor flow-cytometry data, (*31*) analysis also showed that vaccine-specific CD4^+^ T cells from all adult and elderly donors demonstrated similar fundamental characteristics (Fig. S5). However, the elderly induced significantly fewer vaccine-specific CD4^+^ T cells than adults after the first dose, reached the same level as that of adults after the second dose, and again exhibited significantly lower levels at 3 months (Fig. 1A and B). The frequencies of AIM^+^ CD4^+^ T cells before vaccination, which may include naïve, as well as cross-reactive memory phenotype cells (*32, 33*), did not show significant associations with those after vaccination at any time point (Fig. 1C), although a weak correlation was observed after the first dose, which was in agreement with a previous report (*29*). Notably, the cell size according to forward scatter (FSC) of flow cytometry, an indicator of T-cell activation, peaked after the first dose in adults but after the second dose in the elderly and decreased more significantly in the elderly during the contraction phase at 3 months (Fig. 1D).

**Fig. 1.**
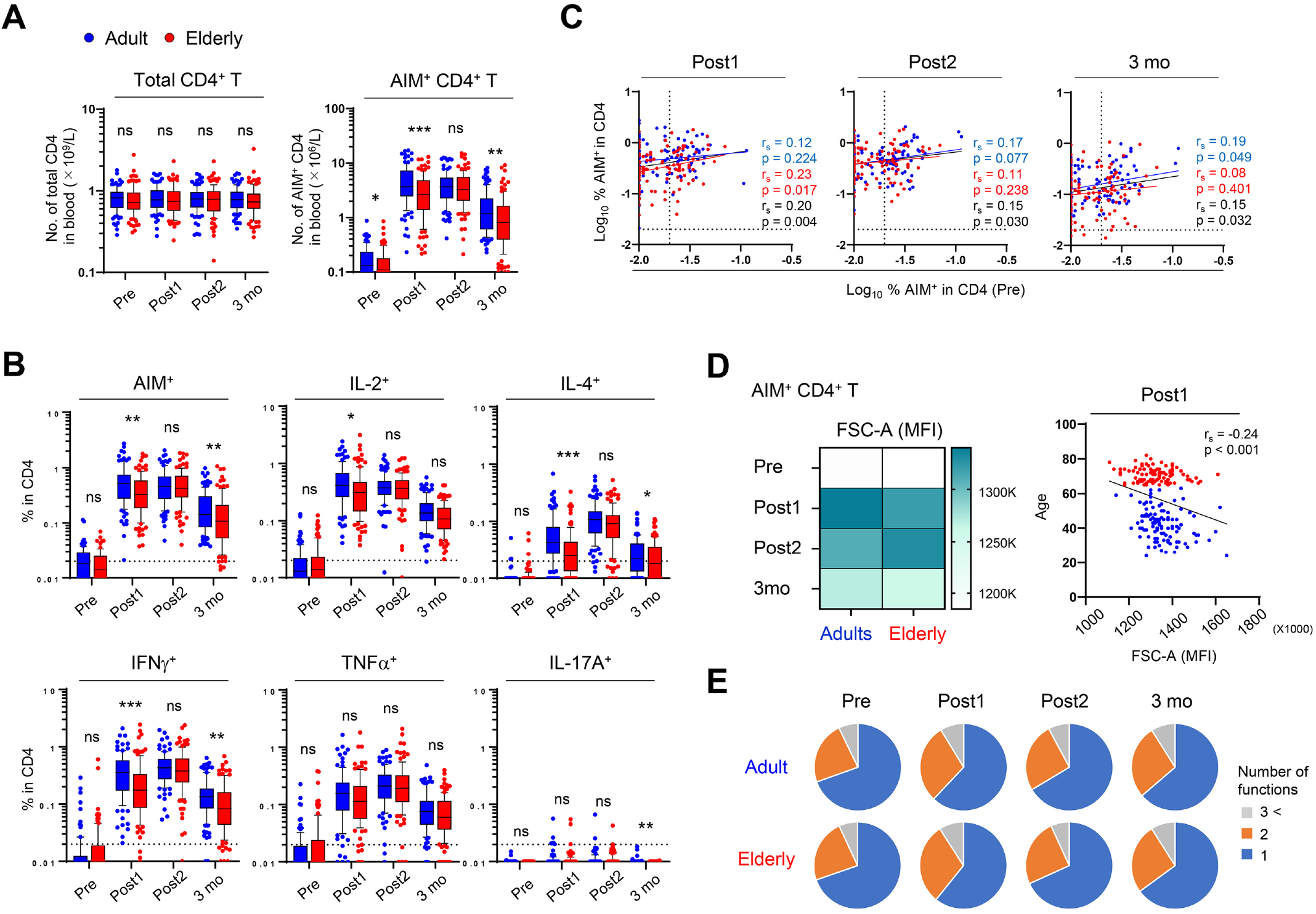
Delayed induction and early contraction of vaccine-specific CD4+ T cells in the elderly. (A) Absolute number of total and AIM^+^(CD137^+^CD154^+^) CD4^+^ T cells in blood. Pre, Postl, Post2, and 3 mo represents the sampling point before vaccination, after the first dose, after the second dose, and 3 months after the first dose, respectively. (B) Frequency of ATM^+^ and cytokine^+^ CD4^+^ T cells. (C) Correlation between the percentage of AIM^+^ CD4^+^T cells before and after vaccination. Percentages of ATM^+^ T cells were transformed into logarithmic values. (D) Heatmap representing the median of mean fluorescent intensity (MFI) of FSC-A in AIM^+^ CD4^+^T cells from adults and the eldely (left). Correlation of MFI of FSC-A in AIM^+^ CD4^+^T cells after the first dose and age of donors (right). (E) Proportions of multiple cytokine-expressing CD4^+^ T cells. The blue, orange, and gray colors in pie charts depict the production of one, two, and more than three cytokines, respectively. (A and B) Box plot represents median with interquartile range (IQR). The whiskers are drawn to the 10th and the 90th percentiles. Points below and above the whiskers are drawn as individual points. (B and C) The dotted line indicates the lower detection limit. Statistical comparisons across cohorts were performed using the Mann-Whitney test. Spearman’s rank correlation (rs) was used to identify relationships between two variables, with a straight line drawn by linear regression analysis. *p < 0.05, **p < 0.01, ***p < 0.001. ns, not significant. Blue, red, and black characters represent the results of statistical test from adults, the elderly, and both groups, respectively.

Major cytokines induced in CD4^+^ T cells after vaccination were interleukin (IL)-2, interferon (IFN)γ, and tumor necrosis factor (TNF)α, whereas IL-4^+^ and IL-17^+^ cells were fewer in frequency in both groups (Fig. 1B). Boolean analysis of cytokine-expressing cells indicated that the two and multiple cytokine-producing cells increased after the first dose and decreased thereafter in both groups (Fig. 1E). However, the frequencies of cytokine^+^ cells, especially IFNγ^+^ cells, in the elderly were significantly lower after the first dose and at 3 months than those in adults (Fig. 1C), which is similar to the kinetics of AIM^+^ cells. Indeed, the negative correlations between age and AIM^+^ or IFNγ^+^ cells after the first dose and at 3 months but not after the second dose were observed in correlation analysis (Fig. S6). Previous studies revealed that cytomegalovirus (CMV) infection and gender differences could affect vaccine responses (*34, 35*). No significant differences were found in the frequencies of AIM^+^ and cytokine^+^ CD4^+^ T cells between male and female or age-matched CMV IgG-seropositive and -seronegative individuals, except for the higher frequency of AIM^+^ cells after the first dose in females (Fig. S7A and B). These results suggested that vaccine-specific CD4^+^ T cells showed similar characteristics in both groups, but that the elderly exhibited delayed induction and rapid contraction after vaccination.

### Delayed induction of vaccine-specific C–X–C motif chemokine receptor 3 (CXCR3)^+^ circulating (c)Tfh cells in the elderly is associated with lower IgG level

We then assessed humoral responses. As reported previously, the mRNA vaccination induced robust IgG responses in all donors, and peak levels of anti-receptor-binding domain (RBD) IgG were observed after the second dose in both groups (*14*) (Fig. 2A). Additionally, IgM response peaked after the second vaccine dose, and we observed strong correlations between IgM and IgG responses after the first and second doses (Fig. 2A and S8A), indicating the simultaneous production of IgM and IgG, as reported previously (*36*), irrespective of age. Although the antibody levels were highly variable among individuals, even within the same age cohort, we observed a negative correlation between age and peak IgG titers after the second dose (r = −0.39; p < 0.001) (Fig. 2B). The peak antibody concentrations in the elderly [median (interquartile range; IQR), 18,200 (5,845) AU/mL] were approximately 40% lower in median as compared with those in adults [median (IQR), 27,200 (7,850) AU/mL] (Fig. 2A). Moreover, the median antibody titers at 3 months decreased to ∼20% of those at the peak and were strongly correlated with those after the second dose (Fig. S8B), suggesting that the gradual decline in antibody levels mostly reflected a natural decay of the antibodies produced at peak response. There was a trend of higher IgG levels in females relative to that of males at every time point, with the values being significantly higher at the pre-vaccination stage and after the second dose (Fig. S8C), which was consistent with the previous report (*37*). However, no significant differences were observed in IgG titers between age-matched CMV-seropositive and -seronegative individuals (Fig. S8D).

**Fig. 2.**
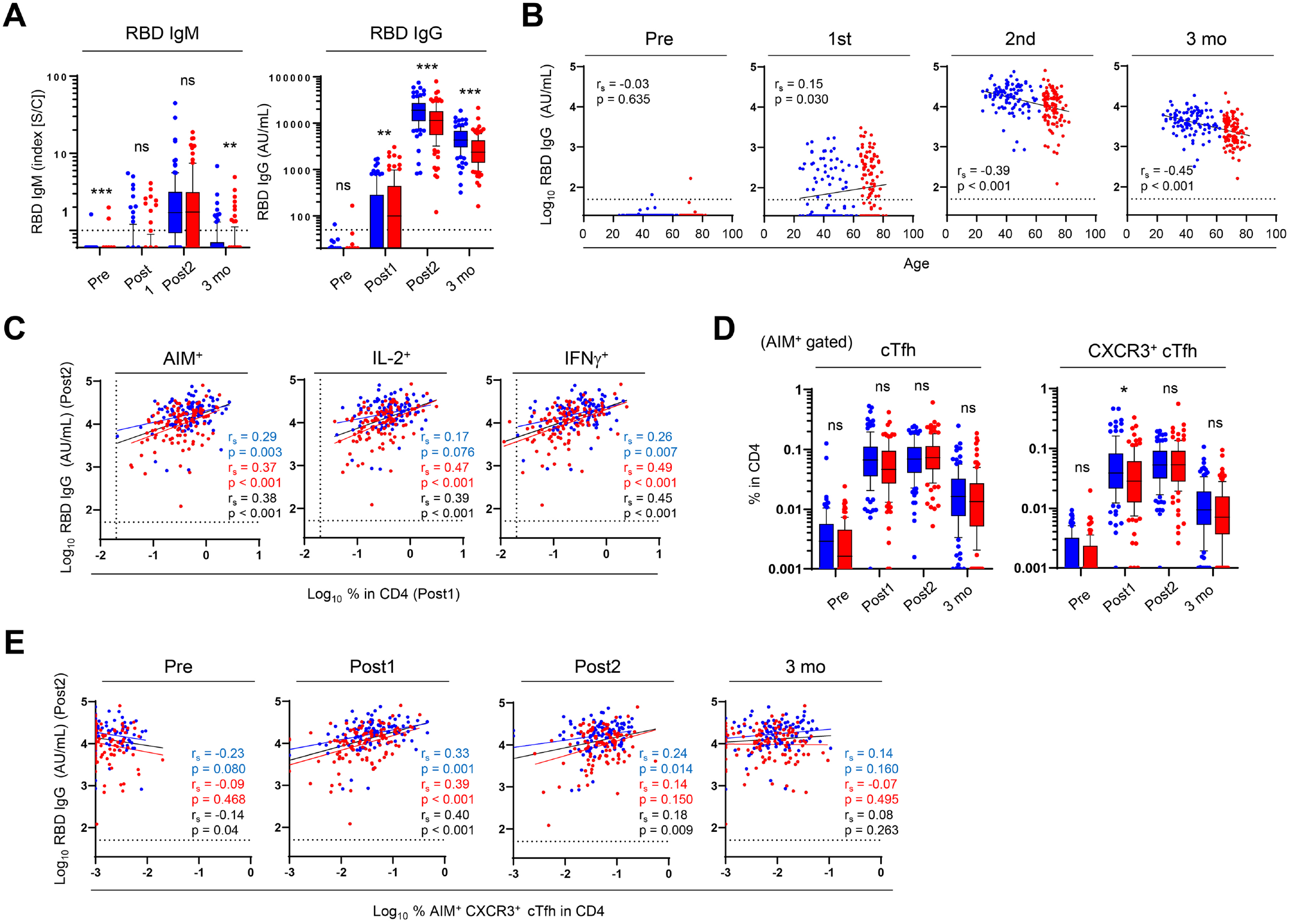
Delayed induction of vaccine-specific CXCR3+ cTfh in the elderly is associated with their lower IgG titer. (A) Concentration of anti-RBD IgM and IgG antibodies. (B) Correlation between the concentration of anti-RBD IgG antibody and age of donors. (C) Correlation between the concentration of anti-RBD IgG antibody after the 2nd dose and the percentage of vaccine-specific CD4^+^T cell after the 1st dose. (D) Frequency of AIM^+^ cTfh and AIM^+^ CXCR3^+^cTfh cells. (E) Correlation between the concentration of anti-RBD IgG antibody after the 2nd dose and the percentage of AIM^+^CXCR3^+^cTfh cells. Box plot represents median with interquartile range (IQR). The whiskers are drawn to the 10th and 90th percentiles. Points below and above the whiskers are drawn as individual points. The dotted line indicates the lower detection limit. Statistical comparisons across cohorts were performed using the Mann-Whitney test. Spearman’ s rank correlation (rs) was used to identify relationships between two variables, with a straight line drawn by linear regression analysis. For correlation analysis, AIM^+^ and cytokine^+^ percentages and concentration of anti-RBD IgG antibody were transformed into logarithmic values. *p < 0.05, **p < 0.01, ***p < 0.001. ns, not significant.

To determine a mechanism for the age-related and individual heterogeneities in antibody responses, we investigated associations between antibody levels and CD4^+^ T-cell responses. The peak IgG concentrations in the elderly showed a strong correlation with AIM^+^, IL-2^+^, or IFNγ^+^ CD4^+^ T cells after the first dose (Fig. 2C), but not after the second dose (Fig. S8E). Tfh cells are a specified subset of CD4^+^ T cells and play key roles in antibody production and germinal center reactions (*38*). The first vaccination rapidly provoked antigen-specific AIM^+^ circulating Tfh (cTfh) cells (CXCR5^+^ CD4^+^ T cells) in peripheral blood (*39*), with no significant differences in the percentages of cTfh cells between the two groups at any given time (Fig. 2D). However, we noted that the levels of CXCR3^+^ cTfh cells (*40-42*) which produce IFNγ and are a major cTfh subset induced by mRNA vaccination (*43*), were significantly lower in the elderly after the first dose (Fig. 2D). Furthermore, we observed a correlation between peak IgG concentration and CXCR3^+^ cTfh cell frequency after the first dose but not at other time points (Fig. 2E). These results strongly suggested that the lower peak IgG level in the elderly was due to the delayed induction of vaccine-specific CD4^+^ T cells, including CXCR3^+^ cTfh cells.

### Fewer systemic AEs after the second dose are relevant to the delayed induction of a vaccine-specific CD4^+^ T-cell response

Although most studies indicate that AEs decrease with age, the associations identified between AEs and antibody responses are largely based on studies involving small cohorts and/or differences in analytical strategies (*21-27*). Furthermore, studies have not addressed whether T-cell responses correlate with AEs. In the present study, to minimize individual differences in the definition of AE severity, we reviewed the medical history of each donor. The frequencies of various systemic AEs (e.g., fever, fatigue, headache, myalgia, arthralgia, and chill) were significantly higher after the second dose than after the first dose (Fig. 3A), which was consistent with previous reports (*14, 44*). Moreover, this confirmed that systemic AEs but not local AEs (pain at an injection site) after the second dose were more commonly observed in adults than the elderly (Fig. 3A). Additionally, the frequency of participants who self-administered antipyretics after the second dose was higher in adults (53.3%) than in the elderly (8.3%), suggesting that AE frequency and degree were significantly underestimated in adults. We then compared the IgG concentration and CD4^+^ T-cell response (AIM^+^ cells) among individuals at each grade of local or systemic AE after the second dose, selecting fever (the most qualitative AE) as the representative systemic AE for statistical analysis. Local pain appeared to show no effect on IgG titer or AIM^+^ CD4^+^ T-cell frequency at both time points (Fig. 3B). By contrast, donors with systemic AE at grade ≥1 (fever of ≥38°C) showed significantly higher IgG titers after the second dose, as well as higher AIM^+^ CD4^+^ T-cell frequencies after the first dose, as compared with those exhibiting fever at grade 0 (Fig. 3B), with these tendencies being observed in both adults and elderly groups (Fig. 3C and Table S2). IL-2 and IFNγ are the major cytokines secreted by CD4^+^ T cells following mRNA vaccination (Fig. 1B) (*18, 29*) and could subsequently induce flu-like symptoms, such as fever or fatigue, upon systemic administration in humans (*45*). Importantly, percentages of CD4^+^ T cells producing IL-2 or IFNγ after the first dose were significantly higher in the systemic AE^+^ group as compared with those in the AE^−^ group, irrespective of age (Fig. 3C). These data indicated that the rapid induction of a vaccine-specific CD4^+^ T-cell response was associated with a higher antibody response and AEs of higher severity after the second vaccination, suggesting a mechanism for the observed lower antibody titers and fewer AEs after the second dose in the elderly.

**Fig. 3.**
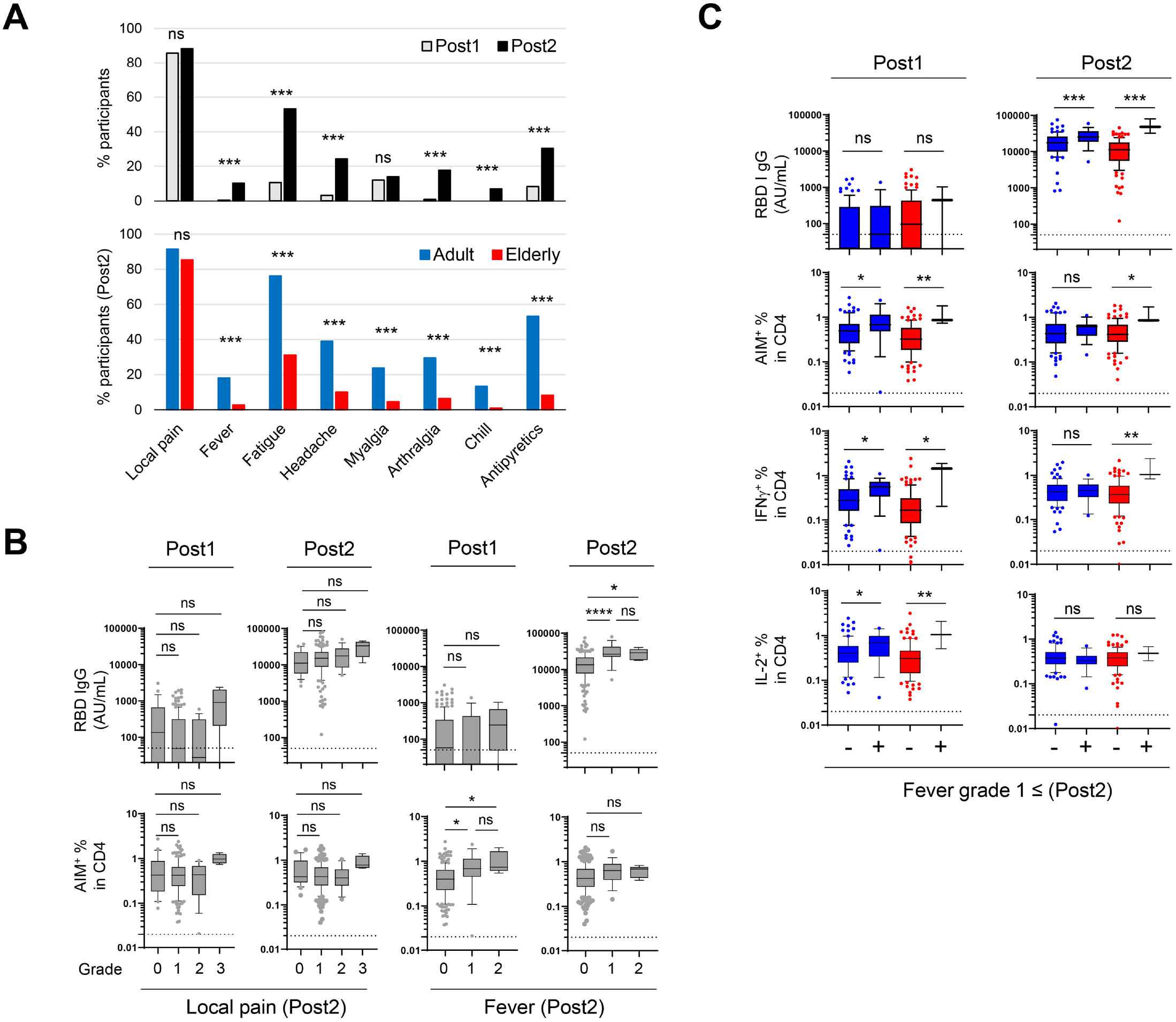
Fewer systemic adverse effects after the second dose are relevant to the delayed induction of vaccine-specific CD4^+^ T cell response. (A) Frequency of donors with adverse effects after vaccination (Upper). Frequency of donors with adverse effects after the second vaccination in adults and elderly (Lower). (B) Concentration of anti-RBD IgG antibody and frequency of AIM^+^ CD4^+^ T cells according to the severity of pain and fever after first and 2nd doses. Multiple comparisons by grade of adverse event symptoms were performed using the Kruskal-Wallis test with Dunn’s *post hoc* test. (C) Concentration of anti-RBD IgG antibody and frequency of AIM^+^ and cytokine^+^CD4^+^T cells according to the emergence of fever after the 2nd dose in adults and elderly. Fisher’s exact test was used to compare the frequency of participants experiencing adverse reactions after vaccination by age group and days from vaccination. A comparison by fever grade in the age group was made using the Mann-Whitney test. Box plot represents median with interquartile range (IQR). The whiskers are drawn to the 10th and 90th percentiles. Points below and above the whiskers are drawn as individual points. The dotted line indicates a lower detection limit. Antipyretics indicate the use of antipyretic medication. *p < 0.05, **p < 0.01, ***p < 0.001. ns, not significant.

### Higher programmed cell death protein-1 (PD-1) expression in vaccine-specific Th1 cells in the elderly correlates with lower CD8^+^ T-cell responses

Similar to AIM^+^ CD4^+^ T cells, AIM^+^ CD8^+^ T-cell induction was significantly delayed in the elderly (Fig. 4A), whereas total CD8^+^ T cells were unchanged during the monitoring period and those in the elderly were significantly lower than in adults, consistent with a previous report (*32*). Th1 cells represent another major CD4^+^ T-cell subset induced by the mRNA vaccine (*19, 29*), with the ability to preferentially produce IFNγ, enhancing CD8^+^ T-cell function, and favorable for antiviral immunity. We confirmed that AIM^+^ CXCR5^−^ Th1 (CXCR3^+^CCR6^−^) cells increased significantly after vaccination, with a peak being observed after the second dose in both groups (Fig. 4B). However, the first vaccination elicited significantly fewer AIM^+^ Th1 cells in the elderly than in adults (Fig. 4B). Consistent with previous reports (*19, 30*), the percentages of AIM^+^ Th1 cells among the total CD4^+^ T cells after the first dose also correlated with the peak antibody titers, with a stronger correlation observed in the elderly (Fig. S9). Importantly, the frequencies of vaccine-specific Th1 cells showed a strong correlation with those of AIM^+^ and IFNγ^+^ CD8^+^ T cells, irrespective of age (Fig. 4C), suggesting a role for Th1 cells in IgG production, as well as CD8^+^ T-cell response.

**Fig. 4.**
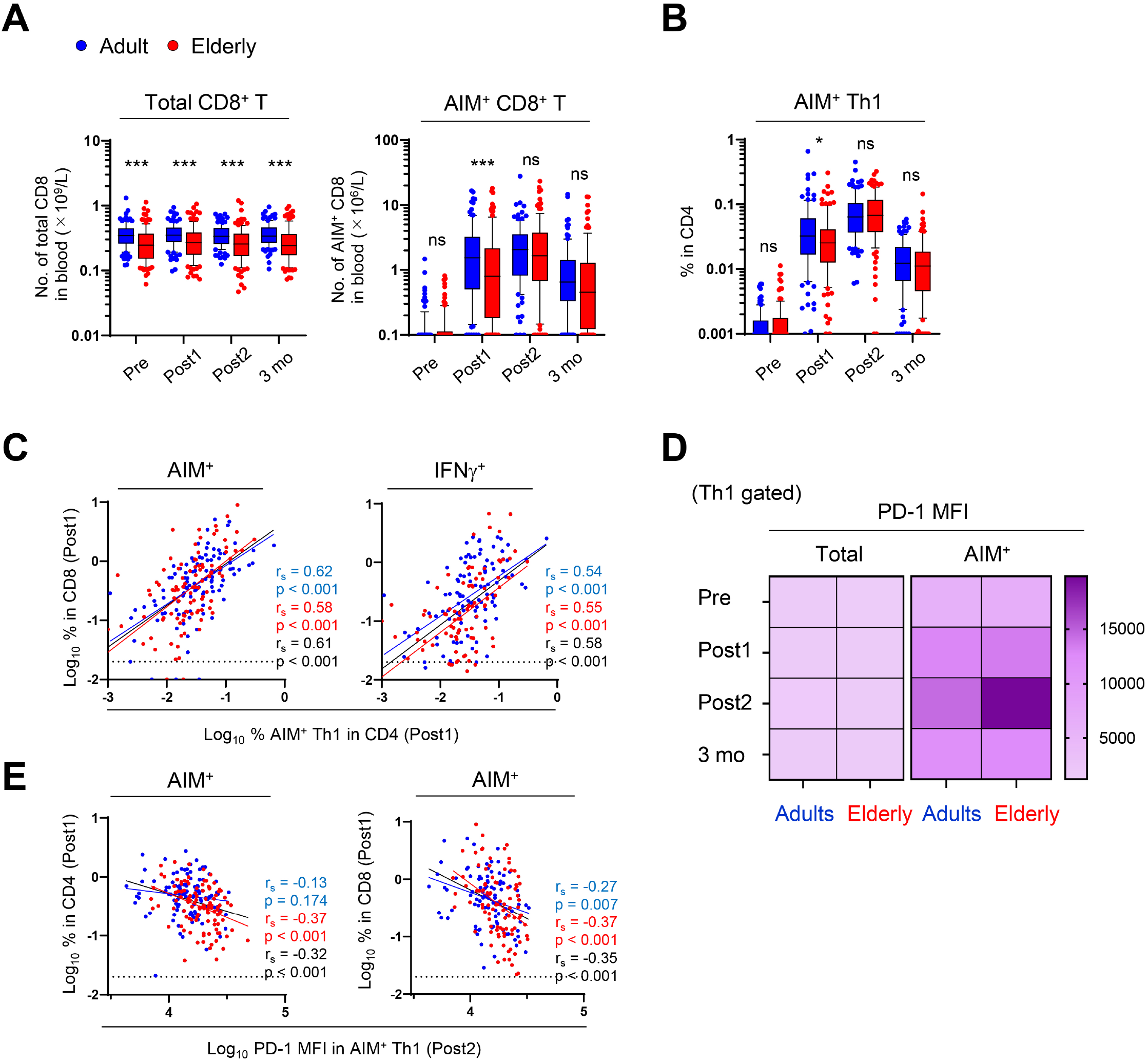
Higher PD-1 expression in vaccine-specific Th1 cells from the elderly is correlated with their lower CD8^+^ T cell responses. (A) Absolute number of total and AIM^+^ (CD69^+^CD137^+^) CD8^+^ T cells in the blood. (B) Frequency of AIM^+^ Th1 cells. (C) Correlation between the percentage of AIM^+^ or IFNγ^+^ CD8^+^T cells and the percentage of AIM^+^ Th1 cells after the 1st dose. (D) Heatmap representing the median of MFI of PD-1 in total and AIM^+^ Th1 cells. (E) Correlation of the percentage of AIM^+^ CD4^+^ and CD8^+^ T cells after the 1st dose and MFI of PD-1 in AIM^+^Th1 cells after the 2nd dose. Samples with at least 0.02% of AIM^+^ CD4^+^T cells were used for the calculation of P0-1 MFI. Box plot represents median with interquartile range (IQR). The whiskers are drawn to the 10th and 90th percentiles. Points below and above the whiskers are drawn as individual points. The dotted line indicates the lower detection limit. Statistical comparisons across cohorts were performed using the Mann-Whitney test. Spearman’s rank correlation (r_s_) was used to identify relationships between two variables, with a straight line drawn by linear regression analysis. Percentages of AIM^+^ and cytokine^+^ T cells were transformed into logarithmic values. *p < 0.05, **p < 0.01, ***p < 0.001. ns, not significant.

To identify a possible mechanism for the low CD4^+^ T-cell response and subsequent compromised humoral and antigen-specific CD8^+^ T-cell responses in the elderly, we focused on PD-1, which is upregulated after T-cell activation and negatively regulate immune responses to prevent excess and/or autoimmune reactions (*46*). PD-1 expression (assessed by mean fluorescence intensity) in total Th1 cells was unchanged in both groups during the observation period (Fig. 4D). However, PD-1 expression in vaccine-specific AIM^+^ Th1 cells gradually increased and peaked after the second dose (Fig. 4D). Importantly, we observed that PD-1 expression after the second dose was significantly higher in the elderly than in adults (Fig. 4D). Furthermore, we observed negative correlations between PD-1 expression and the frequencies of vaccine-specific CD4^+^ and CD8^+^ T cells in the elderly group (Fig. 4E). These results suggested that Th1 cells in the elderly tended to express PD-1 at higher levels upon mRNA vaccination, likely leading to the low humoral and CD8^+^ T-cell responses.

## DISCUSSION

In this study, we investigated age-related differences in antigen-specific T-cell responses induced by SARS-CoV-2 mRNA vaccination in a longitudinal cohort of adult and elderly individuals. We found that CD4^+^ and CD8^+^ T-cell responses in the elderly were lower after the first dose, reached the same level as that in adults after the second dose, but were again significantly lower in the elderly 3 months after vaccination. Thus, the elderly showed delayed induction and early contraction of vaccine-specific T-cells responses.

Previous studies indicate that IgG levels and neutralizing antibody titers decrease with age (*15-17*). However, how the characteristics of T-cell responses in the elderly are associated with the low humoral responses has not been well studied. Consistent with recent studies (*19, 29, 30, 43*), SARS-CoV-2 mRNA vaccination predominantly primed antigen-specific cTfh cells and Th1 cells, and no relative increase in other Th subsets, such as Th2 or Th17, was observed in the elderly. However, we found that, compared with the adults, the elderly showed a significantly lower induction of cTfh cells, especially the CXCR3^+^ cTfh subset, and Th1 cells, both of which correlated with neutralizing antibodies to ancestral virus, as well as variants of concern (*30, 47*). Since the frequencies of both CXCR3^+^ cTfh and Th1 subsets showed positive correlations, the lower induction of one subset was not due to the biased CD4^+^ T-cell responses into either cTfh or Th1 cells but rather the compromised elicitation of Th1-skewed T-cell responses to the mRNA vaccine (*18, 19*). In any case, these data demonstrated that the delayed induction of vaccine-specific CD4^+^ T-cell responses was associated with lower peak IgG titers in the elderly. This may be consistent with a previous study indicating that the rapid induction of antigen-specific CD4^+^ T cells after the first vaccination was associated with coordinated humaral and cellular immunity (*19*), although the cohort included relatively few donors of advanced ages. Indeed, in addition to the age-associated differences, we observed significant individual differences in CD4^+^ T-cell responses after the first dose within the same age cohort of healthy donors, which showed a >10-fold difference in the frequencies of vaccine-specific T cells. Importantly, a poor response to the first dose was also observed in patients harboring solid cancer with lower antibody responses (*48, 49*). These results strongly suggested that CD4^+^ T-cell responses to the first dose is key to the improved consequences of vaccination, and that the elderly tend to have a defect in this process. Moreover, even in the case of SARS-CoV-2 infection, a delay in the development of Tfh cells and subsequent neutralizing antibodies correlates with fatal COVID-19; although, paradoxically, higher IgG production was observed in patients presenting symptoms of higher severity in later stages (*50*). Thus, the delayed induction of CD4^+^ T-cell responses to either vaccination or even viral infection could be a predictive factor for compromised immune competence irrespective of age.

The mechanisms for the delayed induction of CD4^+^ T-cell response remain to be determined. A previous report suggested a correlation between the frequencies of pre-existing SARS-CoV-2 spike-reactive memory CD4^+^ T cells and vaccine-induced CD4^+^ responses after a low first vaccine dose (*29*), whereas the present results indicated no correlation between them at any given time, suggesting little impact of cross-reactive T cells on vaccine responses, at least with the current standard vaccine doses. Additionally, the vaccine-induced CD4^+^ T cells were phenotypically similar in both groups and were mainly and stably mapped to CM subsets after the first dose up to 3 months. However, notably, the cell sizes of vaccine-specific CD4^+^ T cells after the first dose in the elderly were significantly smaller than those in adults, suggesting an inefficient activation of CD4^+^ T cells in the elderly. Furthermore, the frequencies of vaccine-specific CD4^+^ T cells in the elderly reached the same level as those in adults after the second dose but decreased faster, which might reflect an accelerated cell death of effector T cells (*51*). Importantly, the findings of the present study suggested a mechanism for the age-related qualitative difference in CD4^+^ T-cell response: Th1 cells in the elderly tend to express a higher level of PD-1 in response to the mRNA vaccine. Although the CD4^+^ compartment is relatively well maintained qualitatively and quantitatively compared with CD8^+^ T cells, especially in humans (*11, 32*), several T-cell intrinsic defects, such as T-cell receptor desensitization or epigenetic changes, have been reported (*52, 53*). T-cell extrinsic factors, including a defect in antigen-presenting cells with age, were also observed (*54*). Further investigation is warranted to determine which factor primarily contributes to the delayed induction of CD4^+^ T-cells responses to mRNA vaccines. In addition to the age-associated differences, we observed individual variability in T-cell and humoral responses to the mRNA vaccine, as high responders were also found in the elderly population and low responders in the adult group. Therefore, further studies to elucidate whether the low responders in adults show accelerated T cell and/or immune aging will be critical to understanding the mechanisms associated with the large individual variations in immune responses to the mRNA vaccine.

Another key observation was the association between AEs and mRNA-vaccine-induced T-cell responses. Correlations of AEs with antibody titers vary across different studies (*55*), and few studies have analyzed the correlation between AEs and T-cell responses. The present study clearly demonstrated that individuals who exhibited systemic AEs of grade ≥1 (fever of ≥38°C) showed a significantly higher peak of IgG titer and T-cell responses after the first dose compared with those with a fever at grade 0 (fever of <38°), irrespective of age. Thus, it is an intriguing possibility that a high number of effector and memory T cells efficiently induced by the first dose might rapidly produce large amounts of cytokines in response to the second dose. IFNγ^+^ cells were most significantly reduced in CD4^+^ T cells from the elderly group after the first dose. Importantly, IFNγ is a potent inducer of flu-like symptoms, including fever (*45*), and plays an important role in humoral responses (*56*), which likely explains correlations observed among T-cell responses after the first dose, AEs after the second dose, and peak IgG titers. This hypothesis is consistent with a previous study indicating that IFNγ, which is mainly produced by Th1 cells or natural killer cells and not type I IFNs, is the first and primary cytokine demonstrating significant increases at day 1 after the second dose, accompanied by subsequently enhanced levels of phosphorylated signal transducer and activator of transcription (pSTAT)3 and pSTAT1 in multiple cell types and suggestive of a systemic effect of IFNγ (*57*). The group with fever at grade 0 included individuals who showed high levels of T-cell responses and antibody titers, which were much higher than the median of the group with fever at grade ≥1. In the present cohort, ∼30% of participants self-administered antipyretics after the second dose, which might have at least partially affected the results. Altogether, this study strongly suggested the high degree of systemic reactogenicity following delivery of the mRNA vaccine might be an indicator of a strong T cell expansion and Th1 response, which lead to efficient humoral immune responses.

This study has several limitations. We compared the T-cell response and humoral immunity to mRNA vaccines between adults and the elderly. Although individuals ≥65 years of age are commonly defined as the elderly, there is no clear medical or biological evidence to support this definition. Second, vaccination could affect various immune cell subsets other than T cells, and we did not analyze other immune cell types that are critical for vaccine-induced immunity, including antigen-presenting cells and B cells. Moreover, we only investigated the frequencies and cytokine production of vaccine-specific T cells in peripheral blood; therefore, it remains unclear whether other immune cell types or T cells in secondary lymphoid organs, where actual immune responses occur, differ between the two groups and how the difference affects vaccine-induced immune responses. Third, we evaluated only anti-RBD antibody titer but not neutralizing activity, in which cTfh cells play an important role (*38*), although previous studies report that these two parameters are highly correlated (*29, 58*). Finally, we provided evidence only of a correlation between T-cell responses after the first dose with antibody responses in the elderly, as well as AEs irrespective of age. Further studies are needed to investigate causal relationships among these parameters.

In conclusion, we demonstrated the characteristics of immune responses to the mRNA vaccine BNT162b2 in the elderly, revealing a delayed induction and early contraction of antigen-specific CD4^+^ T cells. Specifically, the lower induction of CD4^+^ T cells after the first dose may indicate the lower antibody responses and CD8^+^ T-cell expansion, as well as fewer AEs, in the elderly. This study provides insights for the development of vaccines with higher efficacy and the establishment of a vaccine schedule suitable for the elderly.

## MATERIALS AND METHODS

### Study design

This longitudinal study was reviewed and approved by the Kyoto University Graduate School and Faculty of Medicine, Ethics Committee (R0418). Two hundred and twenty-five participants applied to participate in the study. At the time of enrollment, all donors provided written informed consent, in accordance with the Declaration of Helsinki. Donors were required to be 20 y or older. For the first and second doses, only participants who received Pfizer BNT162T2 were considered eligible. Participants received a BNT162b2 prime dose on day 0 and a boost dose on around day 21, as recommended by the manufacturer. Blood sampling points were set with an allowance (Fig. S1A). They were followed up for medical inquiries at each visit.

All donors were otherwise healthy and did not report any ongoing severe medical conditions, including cancer, gastrointestinal, liver, kidney, cardiovascular, hematologic, or endocrine diseases. Participants taking medications that may affect the immune system, including steroids or immunomodulatory drugs, were excluded. Blood samples were collected at the Ki-CONNECT and Clinical BioResource Center (CBRC) at Kyoto University Hospital. Samples were de-identified using an anonymous code assigned to each sample. Only samples without bloodborne pathogens, including HIV, HTLV-1, HBV, and HCV, were used for subsequent experiments. Six did not meet the eligibility criteria, and a total of 219 individuals consisting of 107 adults (aged less than 65 years, mainly workers in Kyoto University Hospital) and 112 elderly individuals (aged more than 65 years, mainly ordinary citizens) were enrolled in the study. Two patients were lost to follow-up, and one was stopped because of mRNA-1273 injection as the primary and boost vaccination (Fig. S1B). Their characteristics, including age, sex, and serology, are summarized in Table 1.

### PBMCs isolation, cryopreservation, and thawing

Whole blood was drawn into Vacutainer CPT™ Cell Preparation Tubes with sodium citrate (BD biosciences), according to the manufacturer’s instructions, and processed within 2 hours to isolate peripheral blood mononuclear cells (PBMCs). Isolated PBMCs were resuspended in CELLBANKER 1 (ZENOGEN PHARMA) at a concentration of 8×106 cells/mL and aliquoted in 250 or 500 ml per cryotube. Samples were stored at −80 ℃ on the day of collection and in liquid nitrogen until used for the assays. Cryopreserved PBMCs were thawed in pre-warmed X-VIVO15 (LONZA) without serum. After centrifugation, the cells were washed once and used directly for assays, as described below.

### Complete blood counts

Whole blood was collected in ethylenediaminetetraacetic-2Na tubes. The analysis was performed using an Automated Hematology Analyzer XN-9000 (Sysmex Corporation) at the Department of Clinical Laboratory, Kyoto University Hospital.

### Serology

Whole blood was collected in a Venoject VP-P075K (Terumo) blood collection vessel for serum isolation. The serum separator tubes were centrifuged for 4 min at 1100 *g* at 4 ℃. The serum was then removed from the upper portion of the tube, aliquoted, and stored at −80℃.

Anti-SARS-CoV-2 (N protein) IgM/IgG levels in the serum were measured using an Elecsys Anti-SARS-CoV-2 with cobas8000 (Roche Diagnostics KK) at the Department of Clinical Laboratory, Kyoto University Hospital. Anti-SARS-CoV-2 RBD IgM and IgG levels were measured at LSI Medience (Tokyo, Japan) using ARCHITECT SARS-CoV-2 IgM and ARCHITECT SARS-CoV-2 IgG II Quant (Abbott), respectively. Anti-CMV IgG levels were measured using a chemiluminescence immunoassay (CLIA) at LSI Medience (Tokyo, Japan). The cut-off values for Anti-SARS-CoV-2 (N protein) IgM/IgG, Anti-SARS-CoV-2 RBD IgM, IgG and Anti-CMV IgG were 1.0 cutoff index (COI), 1.0 (COI), 50 (AU/mL) and 6.0 (AU/mL), respectively.

### Peptide pools

PepTivator® SARS-CoV-2 Prot_S Complete peptide pools (Miltenyi Biotech) were diluted in distilled water (DW) and used for vaccine-specific T-cell stimulation. The S peptide pool contains 15-mer peptides that overlap by 11 amino acids and cover the complete protein-coding sequence (aa 5–1273) of the surface or spike glycoprotein (S) of SARS coronavirus 2 (GenBank MN908947.3, Protein QHD43416.1). Peptide pools were added to the culture medium at a final concentration of 0.6 nmol/mL.

### Activation-induced marker (AIM) assay

PBMCs were cultured in 100 µL of X-VIVO15 medium supplemented with 5% human AB serum for 23 h at 37 ℃ in the presence of SARS-CoV-2 peptide pools (0.6 nmol/mL) and CD40 blocking antibody (0.5 µg/mL, Miltenyi Biotech) in 96-well U-bottom plates at 1×10^6^ PBMCs per well. An equal volume of DW was used as a negative control. After stimulation, cells were stained with fluorochrome-conjugated surface antibodies at pre-titrated concentrations in the presence of FcR blocking (Miltenyi Biotech) for 20 min at 4 ℃. The cells were then washed and stained with Ghost Dye™ Red 710 (TONBO) to discriminate between viable and non-viable cells. After the final wash, the cells were resuspended in 100 µL PBS with 2% FBS (FACS buffer) for flow cytometry. The antibodies used in the AIM assay are listed in Table S1. Vaccine-specific AIM^+^ T cells were defined based on the co-expression of CD154 (CD40L) and CD137 for CD4^+^ T cells and CD137 and CD69 for CD8^+^ T cells (Fig. S2B). Vaccine-specific responses were quantified as the frequency of AIM^+^ cells in stimulated samples, with background subtraction from paired DW controls (Fig. S2B).

### Intracellular cytokine staining (ICS) assay

As similar to the AIM assay, PBMCs were cultured in 100 µL of X-VIVO15 medium supplemented with 5% human AB serum for 23 h at 37 ℃ in the presence of SARS-CoV-2 peptide pools (0.6 nmol/mL) at 1×10^6^ PBMCs per well. An equal volume of DW was used as the negative control. Four hours before ICS administration, brefeldin A (BioLegend) (1:1000 dilution) was added to the medium. After stimulation, cells were stained with fluorochrome-conjugated surface antibodies at pre-titrated concentrations in the presence of FcR blocking for 20 min at 4 ℃. Cells were then washed and stained with Ghost Dye™ Red 710 to discriminate between viable and non-viable cells. The stained cells were then fixed with IC fixation buffer (Thermo Fisher Scientific) for 30 min at 4 ℃ and washed twice with permeabilization buffer (Thermo Fisher Scientific) and subsequently stained for intracellular IL-2, IL-4, IL-17A, IFNγ, TNFα, perforin, and granzyme (1:100 dilution each) for 30 min at room temperature. After the final wash, the cells were resuspended in 100 µL of FACS buffer for flow cytometry. The antibodies used in ICS assays are listed in Table S1. Vaccine-induced cytokine-producing T cells were quantified as the frequency of cytokine^+^ cells in stimulated samples, with background subtraction from paired DW controls (Fig. S3B).

### Flow cytometry and FCS data analysis

All AIM and ICS assay samples were acquired using Northern Light 3000 (Cytek). FCS 3.0 data files were exported and analyzed using FlowJo software version 10.8.1. The detailed gating strategies for individual markers are described in Fig. S2 and S3. The subset definitions and gating strategies are outlined in the text or figure legends. The absolute number of a defined subset of CD4^+^ and CD8^+^ T cells was obtained by multiplying the number of CD4^+^ or CD8^+^ T cells by the percentage of the corresponding subsets. To calculate the frequency of each subset, samples with at least 0.02% AIM^+^ CD4^+^ T cells are indicated in the figures because a very low frequency of AIM^+^ cells influence the results. Proportions of multiple cytokine-expressing CD4^+^ T cells (IFNγ, IL-2, IL-4, IL-17A, and TNFα) were assessed by Boolean analysis as reported previously (29). All the background-subtracted subpopulations with at least 0.005% of cytokine^+^ cells were combined into a sum of cytokine-expressing CD4^+^ T cells based on the number of functions per group. Samples with a summed frequency higher than 0.01% were considered for the analysis. Relative proportions of number of functions were displayed as a pie chart.

### opt-SNE and FlowSOM

Dimensionality reduction of multicolor flow cytometry data obtained from AIM assays was performed using OMIQ software. FCS 3.0 data from all donors were imported. Up to 40 AIM^+^ CD4^+^ T cells were sub-sampled and merged for analysis. The subsampling counts were derived from the 25 percentile in the corresponding subset, which could pool AIM^+^ cells evenly from most donors. The markers applied to the opt-SNE and FlowSOM are described in the figure legends. The parameters used were as follows: opt-SNE: max iterations = 1000, opt-SNE end = 5000, perplexity = 30, theta = 0.5, components = 2, random seed = 6925, verbosity = 25; FlowSOM: xdim = 10, ydim = 10, rien =10, comma-separated k values = 20, 25, and random seed = 6793.

### Statistical analyses

Statistical analyses were performed using GraphPad Prism 9.0. Statistical details, such as groups, statistical tests, and significance values of the results are provided in the respective figure legends. Two-group comparisons were conducted using a Mann-Whitney test and comparisons between three or more groups were performed using a Kruskal-Wallis test.

## List of Supplementary Materials

Results

Fig S1 to S9

Table S1 and S2

## Acknowledgments

We thank all donors and healthcare personnel involved in this work. We are grateful to Dr. K. Kometani for helpful comments; E. Yamaguchi, C. Takashima, I. Kasumoto, and T. Ikari for technical assistance; S. Takakura, J. Ueda, and A. Yonemura for participant arrangement; other members of our laboratory and hospital staff for assistance; Drs. T. Honjo and E. Hara for AMED project administration; Dr. S. Yamanaka for organizing CiRA Fight Corona Project; and Dr. N. Minato for helpful discussions.

## Funding

This work was supported by the Japan Agency for Medical Research and Development (grant numbers JP21gm5010005 and JP20fk0108454) (to YH), iPS Cell Research Fund (to YH), the COVID-19 Private Fund to the Shinya Yamanaka Laboratory, CiRA, Kyoto University (to YH), Japanese Ministry of Education, Culture, Sports, Science and Technology (MEXT)/Japan Society for the Promotion of Science (grant number 19K23862) (to NJ), Kansai Economic Federation (KANKEIREN) (to YH), and Takeda Science Foundation (to YH).

## Author contributions

Performance of experiments and data analyzes: NJ

Statistical analyses: YH and SM

Organization of the vaccine cohort and medical inquiry: OK, MF, TS, MA, TN, and MM.

Performance of blood tests: MY and MN.

Experiment design: NJ and YH.

Manuscript preparation: NJ and YH.

Conception of the study idea and project mangement: YH

All authors contributed intellectually and approved the manuscript.

## Competing interests

The authors declare that the research was conducted in the absence of any commercial or financial relationships that could be construed as potential conflicts of interest.

## Data and materials availability

All data associated with this study are present in the paper or the Supplementary Materials.

## Notes

### Competing Interest Statement

The authors have declared no competing interest.

