## Supplementary material for "Delayed antigen-specific CD4^+^ T-cell induction correlates with impaired immune responses to SARS-COV-2 mRNA vaccination in the elderly": Supplentary Fig and table

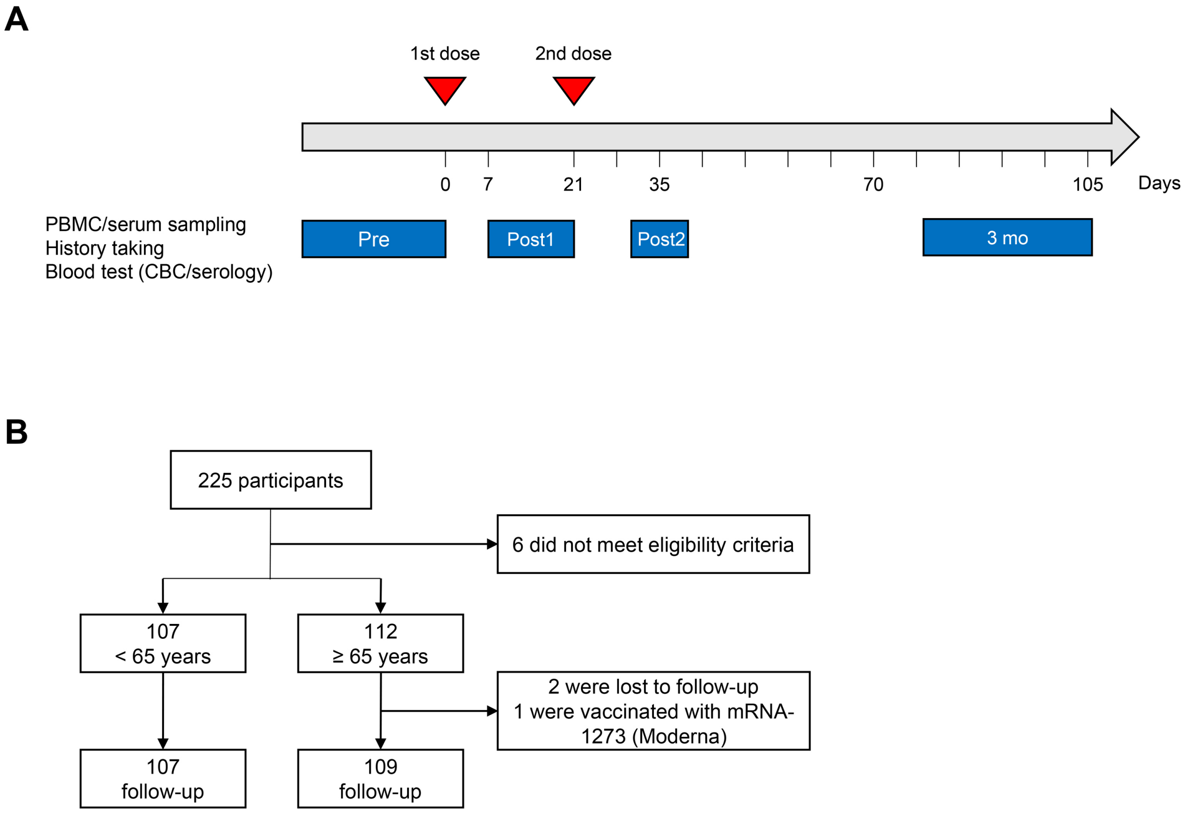


**Fig. S1. Study design.** (A) The participants received a BNT162b2 1st dose on day 0 and a 2nd dose on around day 21. The sampling points were set with an allowance: 7-21 days after the first dose (Post1), 31-39 days after the second dose (Post2), and 79-104 days at the point of 3 months after the first dose (3 mo). The actual vaccinated and sampling days are described in the Results section. CBC; complete blood count. (B) The diagram represents all participants throughout the study.

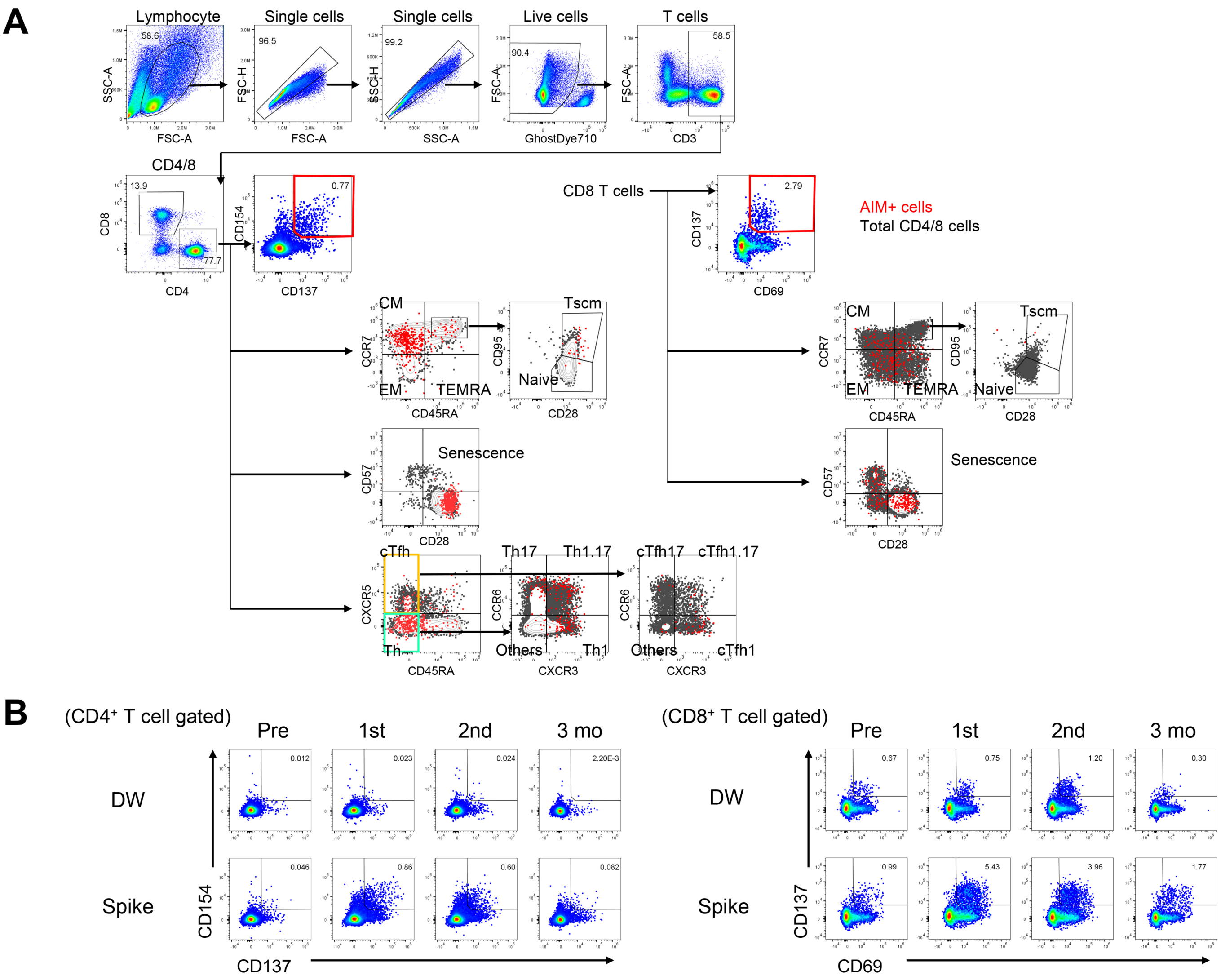


**Fig. S2. Identification and phenotyping of AIM$^+$ T cells.** (A) Representative gating strategy to detect and characterize vaccine-specific T cells in the AIM assay. Briefly, lymphocytes were gated out of all events, and doublets were excluded. Live cells were gated as Ghost Dye$^{TM}$ Red 710$^-$. T cells were then gated as CD3$^+$ and subdivided into CD4$^+$ and CD8$^+$ populations. CD4$^+$ and CD8$^+$ T cells were further characterized for the phenotype such as differentiation, senescence, Th, and cTfh. Differentiation status were defined as naive (CD45RA$^+$CCR7$^+$CD28$^+$CD95$^-$), stem cell memory (CD45RA$^+$CCR7$^+$CD28$^+$CD95$^+$), central memory (CD45RA-CCR7$^+$), effector memory (CD45RA$^-$CCR7$^-$) or terminally differentiated effector memory cells re-expressing CD45RA (TEMRA, CD45RA$^+$CCR7$^-$). T cell senescence was assessed using CD28 and CD57. Th and cTfh were gated as CD45RA$^-$CXCR5$^-$ and CD45RA$^-$CXCR5$^+$, respectively. Th and cTfh were further divided into type 1, 2, 17, and 1/17 subtypes based on the expression of CXCR3 and CCR6. (B) Representative FCM plots displaying AIM$^+$ (CD137$^+$CD154$^+$ for CD4, and CD69$^+$CD137$^+$ for CD8$^+$) T cells after stimulation with the negative control (DW) or spike protein overlapping peptides (spike). Numbers indicate the population percentages in the gates.

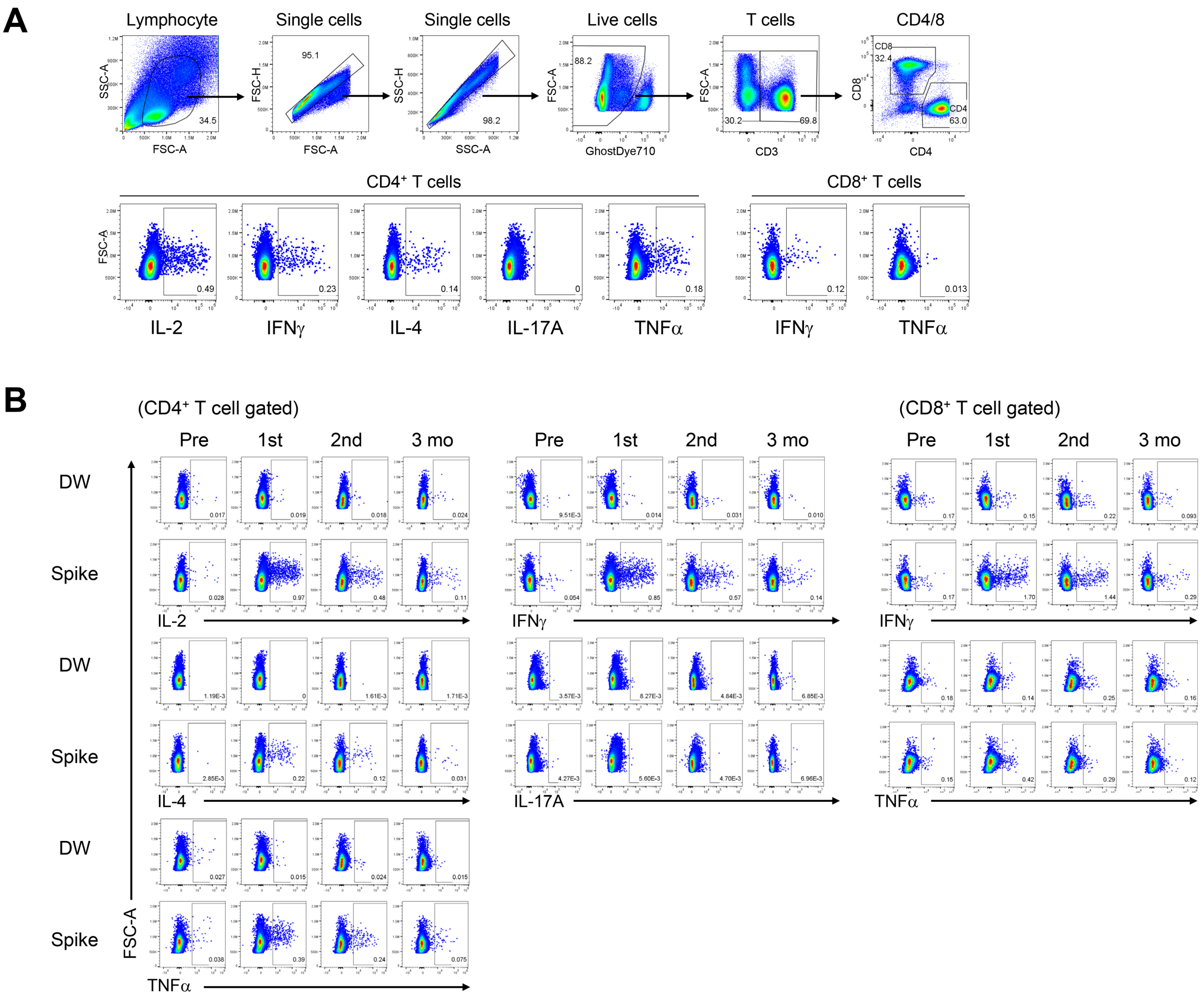


**Fig. S3. Identification of cytokine$^{+}$ T cells.** (A) Representative gating strategy to detect and characterize vaccine-specific T cells in the ICS assay. Briefly, lymphocytes were gated out of all events, and doublets were excluded. Live cells were gated as Ghost Dye$^{TM}$ Red 710$^{-}$. T cells were then gated as CD3$^{+}$ and subdivided into CD4$^{+}$ and CD8$^{+}$ populations. CD4$^{+}$ and CD8$^{+}$ T cells were further analyzed for corresponding cytokines. (B) Representative FCM plots displaying cytokine$^{+}$ CD4$^{+}$ and CD8$^{+}$ T cells after stimulation with the negative control (DW) or spike protein overlapping peptides (spike). Numbers indicate the population percentages in the gates.

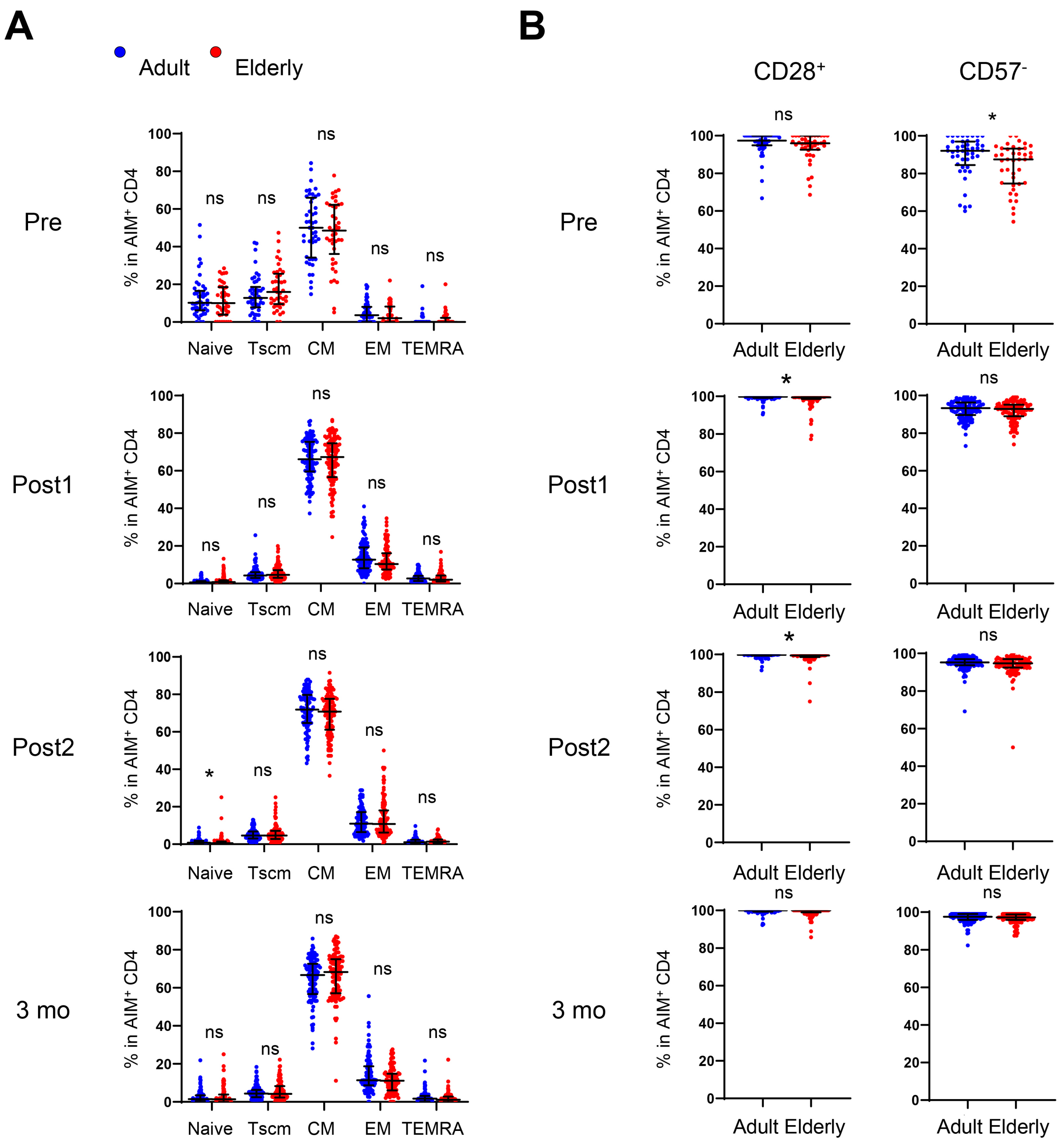


**Fig. S4. Differentiation and senescence status of vaccine-specific CD4⁺ T cells.** (A) Frequency of naive, Tscm, CM, EM, and TEMRA in AIM⁺ CD4⁺ T cells. (B) Frequency of CD28⁺ and CD57⁻ in AIM⁺ CD4⁺ T cells. Samples with at least 0.02% of AIM+ CD4⁺ T cells were analyzed. The centerline and error bars indicate the median ± IQR. Statistical comparisons across cohorts were performed using the Mann-Whitney test. *$p < 0.05$, ns, not significant.

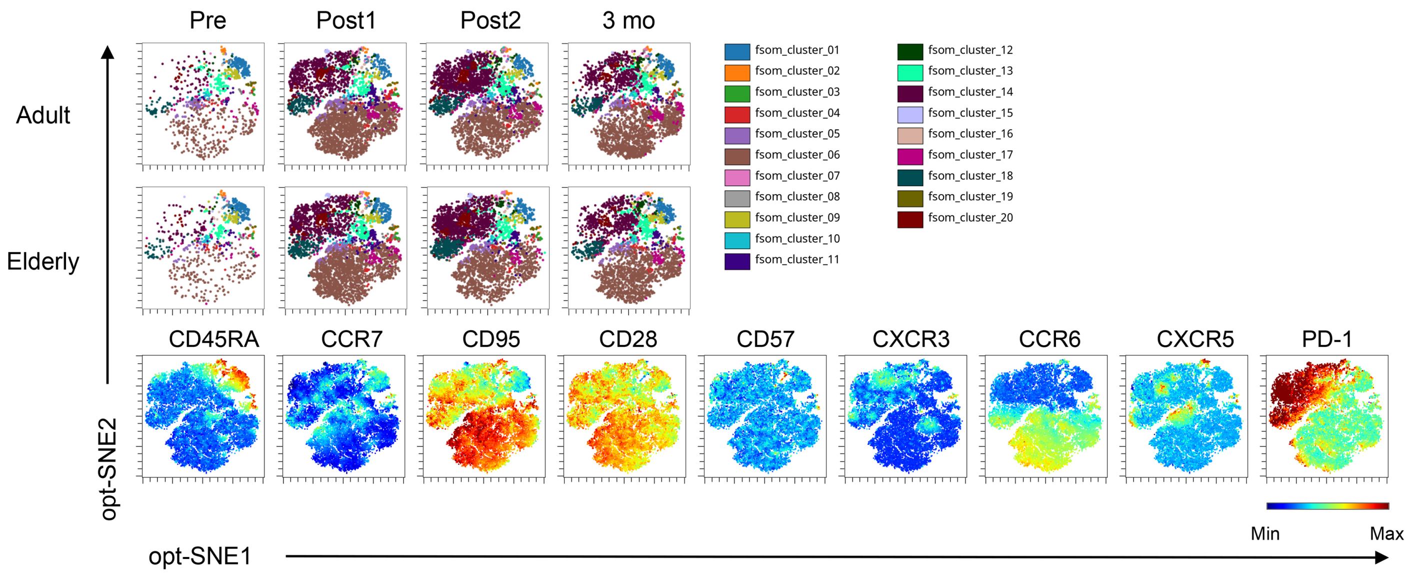


**Fig. S5. opt-SNE for vaccine-specific CD4$^+$ T cells.** FlowSOM clustering of AIM$^+$ CD4$^+$ T cells pooled from adults and the elderly is displayed on opt-SNE plots. The markers described in the bottom row (CD45RA, CCR7, CD95, CD28, CD57, CXCR3, CCR6, CXCR5, and PD-1) were used for the analysis.

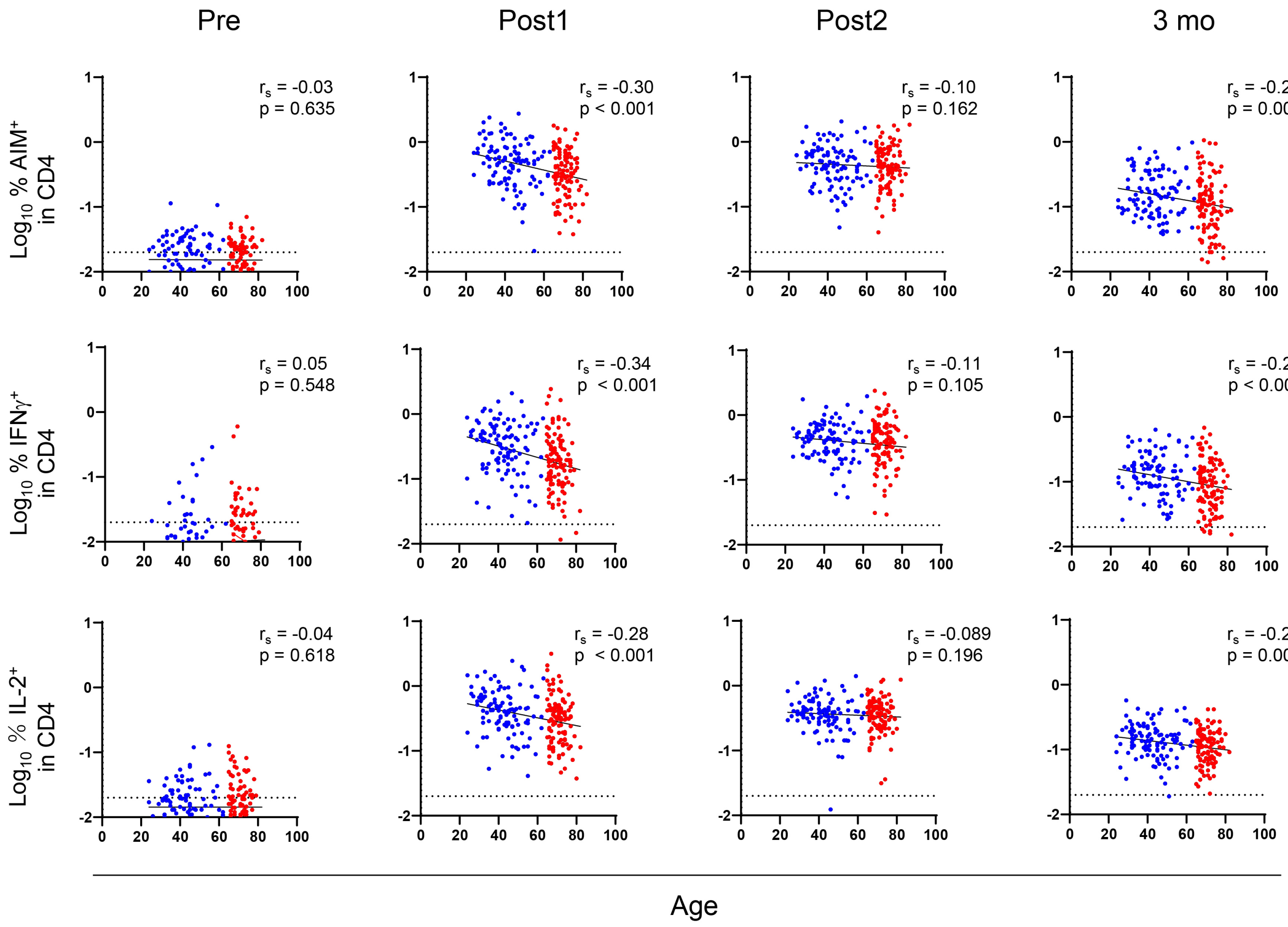


**Fig. S6. Correlation between the frequencies of vaccine-specific $CD4^+$ T cells and age** Correlation between the percentages of $AIM^+$ and $cytokine^+$ $CD4^+$ T cells and donor age. Spearman's rank correlation ($r_s$) was used to identify relationships between two variables, with a straight line drawn by linear regression analysis.

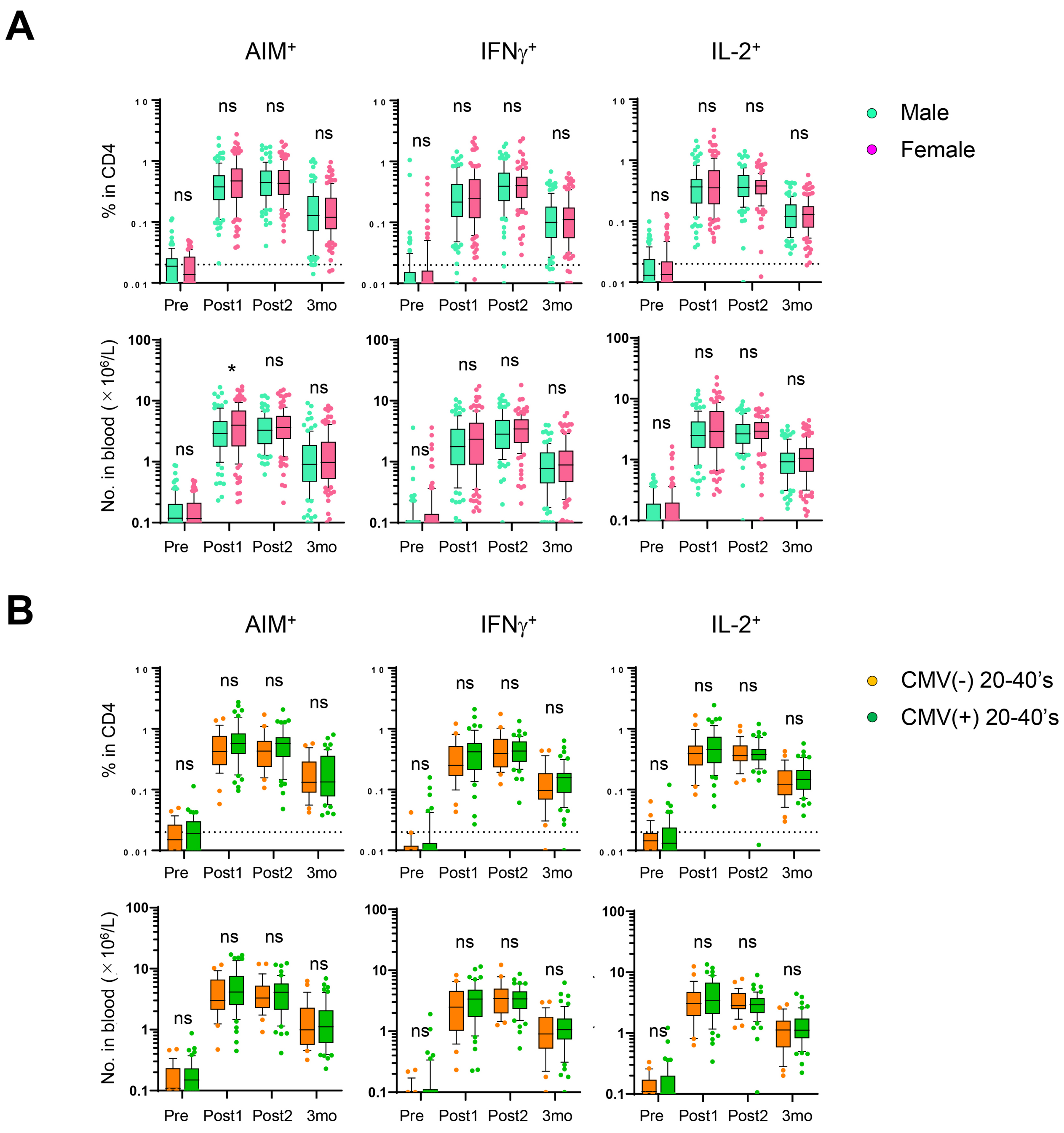


**Fig. S7. Comparison of vaccine-specific CD4$^+$ T cells between male and female, and CMV-seronegative and seropositive.** Frequency and absolute number of AIM$^+$ and cytokine$^+$ CD4$^+$ T cells from male and female donors (A) and CMV-seronegative and -seropositive donors in their 20 to 40s (B). Box plots represent the median and interquartile range (IQR). Whiskers were drawn to the 10th and 90th percentiles. The points below and above the whiskers are drawn as individual points. The dotted line indicates the lower detection limit. Statistical comparisons across cohorts were performed using the Mann-Whitney U test. *$p < 0.05$; ns, not significant.

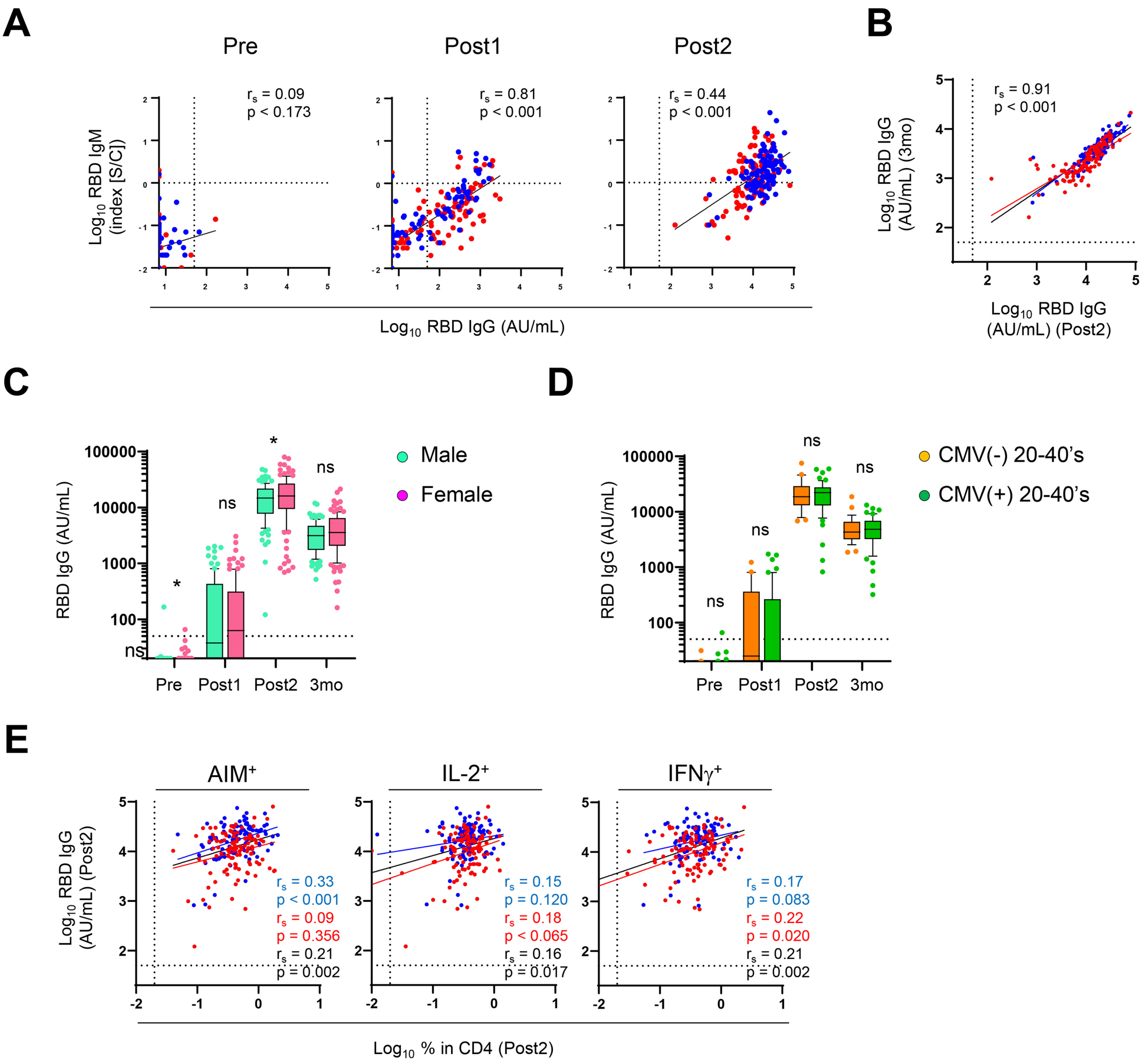


**Fig. S8. Antibody levels and CD4+ T cell responses after vaccination.** (A) Correlation between the concentrations of anti-RBD IgM and anti-RBD IgG antibody at the same sampling points. (B) Correlation between the concentrations of anti-RBD IgG antibody after the 2nd dose and at 3 months. (C, D) Concentration of anti-RBD IgG antibody from male and female donors (C) and CMV-seronegative and seropositive donors in their 20 to 40s (D). (E) Correlation between the concentration of anti-RBD IgG antibody after the 2nd dose and the percentage of vaccine-specific CD4+ T cell after the 2nd dose. Box plot represents median with interquartile range (IQR). The whiskers are drawn to the 10th and 90th percentiles. Points below and above the whiskers are drawn as individual points. The dotted line indicates the lower detection limit. Statistical comparisons across cohorts were performed using the Mann-Whitney test. Spearman's rank correlation ($r_s$) was used to identify relationships between two variables, with a straight line drawn by linear regression analysis. For correlation analysis, AIM+ and cytokine+ percentages and concentration of anti-RBD IgM and IgG antibodies were transformed into logarithmic values. *p < 0.05, **p < 0.01, ***p < 0.001. ns, not significant.

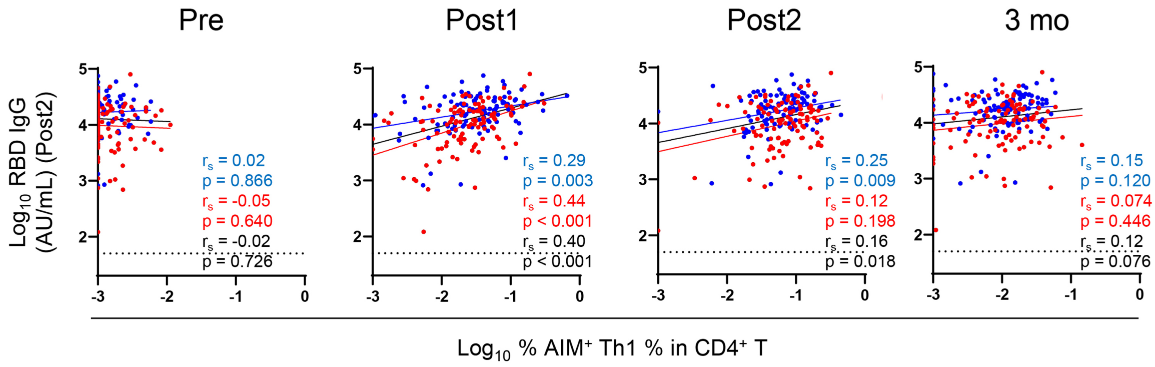


**Fig. S9. Correlation between anti-RBD IgG and Th1 cells.** Correlation between the concentration of anti-RBD IgG antibody after the 2nd dose and the percentage of vaccine-specific Th1 cells. The dotted line indicates the lower detection limit. Spearman's rank correlation ($r_s$) was used to identify relationships between two variables, with a straight line drawn by linear regression analysis. $AIM^+$ percentages and concentrations of anti-RBD IgG antibodies were logarithmically transformed.

AIM

| Marker | Fluorochrome | Clone | Manufacturer | Dilution |
|---|---|---|---|---|
| PD-1 | BV421 | EH12.2H7 | Biolegend | 1:100 |
| CD57 | BV510 | QA17A04 | Biolegend | 1:2500 |
| CD8 | BV570 | RPA-T8 | Biolegend | 1:500 |
| CD154 | BV605 | 24-31 | Biolegend | 1:100 |
| CD69 | BV650 | FN50 | Biolegend | 1:100 |
| CD28 | BV750 | CD28.2 | Biolegend | 1:50 |
| CD95 | BV785 | DX2 | Biolegend | 1:200 |
| CCR6 | FITC | G034E3 | Biolegend | 1:100 |
| CD4 | AF532 | RPA-T4 | Invitrogen | 1:100 |
| CXCR5 | PE | J252D4 | Biolegend | 1:100 |
| CD45RA | PerCP-Cy5.5 | HI100 | Biolegend | 1:100 |
| CD3 | PerCP-eF710 | OKT3 | Invitrogen | 1:250 |
| CD137 | PE/Cy7 | 4B4-1 | Biolegend | 1:100 |
| CXCR3 | APC | G025H7 | Biolegend | 1:50 |
| GD710 | AF700 | - | TONBO | 1:1000 |
| CCR7 | APC-Cy7 | G043H7 | Biolegend | 1:100 |

ICS

| Marker | Fluorochrome | Clone | Manufacturer | Dilution |
|---|---|---|---|---|
| Perforin | BV421 | dG9 | Biolegend | 1:100 |
| CD57 | BV510 | QA17A04 | Biolegend | 1:2500 |
| CD8 | BV570 | RPA-T8 | Biolegend | 1:500 |
| CD45RA | BV605 | HI100 | Biolegend | 1:100 |
| TNFa | BV650 | MAb11 | Biolegend | 1:100 |
| CD28 | BV750 | CD28.2 | Biolegend | 1:50 |
| CD95 | BV785 | DX2 | Biolegend | 1:200 |
| CCR7 | FITC | G043H7 | Biolegend | 1:100 |
| CD4 | AF532 | RPA-T4 | Invitrogen | 1:100 |
| IL-4 | PE | MP4-25D2 | Biolegend | 1:100 |
| Granzyme | PerCP-Cy5.5 | QA18A28 | Biolegend | 1:100 |
| CD3 | PerCP-eF710 | OKT3 | Invitrogen | 1:250 |
| IL-2 | PE/Cy7 | MQ1-17H12 | Biolegend | 1:100 |
| IFNɣ | APC | 4S.B3 | Biolegend | 1:100 |
| GD710 | AF700 | - | TONBO | 1:1000 |
| IL-17A | APC-Cy7 | BL168 | Biolegend | 1:100 |

**Table S1. Antibody list used for AIM and ICS assay.**

Post 1

| Parameters | Fever (Grade) | < 65 years N = 107 (Median ± IQR) | ≥ 65 years N = 109 (Median ± IQR) |
|---|---|---|---|
| RBD IgG | 0 | 14.1 ± 280 | 96.3 ± 419.2 |
| | 1 | 10.95 ± 462 | 225.4 ± 437.2 |
| | 2 | 166.8 ± 252.5 | 1030 |
| AIM⁺ CD4 (%) | 0 | 0.49 ± 0.45 | 0.33 ± 0.39 |
| | 1 | 0.60 ± 0.71 | 1.33 ± 0.93 |
| | 2 | 1.00 ± 1.26 | 0.74 |
| IFNγ⁺ CD4 (%) | 0 | 0.28 ± 0.34 | 0.17 ± 0.24 |
| | 1 | 0.58 ± 0.44 | 1.70 ± 0.40 |
| | 2 | 0.44 ± 0.38 | 0.2 |
| IL-2⁺ CD4 (%) | 0 | 0.41 ± 0.33 | 0.31 ± 0.31 |
| | 1 | 0.69 ± 0.71 | 1.6 ± 1.10 |
| | 2 | 0.60 ± 0.93 | 0.51 |

Post 2

| Parameters | Fever (Grade) | < 65 years N = 107 (Median ± IQR) | ≥ 65 years N = 109 (Median ± IQR) |
|---|---|---|---|
| RBD IgG | 0 | 17550 ± 15862 | 11200 ± 12195 |
| | 1 | 25100 ± 14000 | 64000 ± 32000 |
| | 2 | 23800 ± 19500 | 32200 |
| AIM⁺ CD4 (%) | 0 | 0.43 ± 0.45 | 0.42 ± 0.4 |
| | 1 | 0.63 ± 0.32 | 1.29 ± 0.86 |
| | 2 | 0.59 ± 0.28 | 0.83 |
| IFNγ⁺ CD4 (%) | 0 | 0.43 ± 0.35 | 0.37 ± 0.36 |
| | 1 | 0.54 ± 0.34 | 1.6 ± 1.57 |
| | 2 | 0.35 ± 0.17 | 1 |
| IL-2⁺ CD4 (%) | 0 | 0.37 ± 0.23 | 0.38 ± 0.26 |
| | 1 | 0.33 ± 0.12 | 0.58 ± 0.2 |
| | 2 | 0.34 ± 0.41 | 0.33 |

**Table S2. Antibody levels and $CD4^+$ T-cell responses of individuals by subgrouping them according to the grade of fever**
